# Lipid production from *Zygosaccharomyces siamensis* AP1 using glycerol as a carbon source

**DOI:** 10.1101/2021.03.03.433823

**Authors:** M Ilmi, M Siswantoro

**Affiliations:** Faculty of Biology, Universitas Gadjah Mada, Jl. Teknika Selatan, Sekip Utara, Yogyakarta, 55281

**Keywords:** C/N ratio, cheap carbon source, oleaginous yeast, production profile, wild honey

## Abstract

Yeasts are considered as potential lipid producer because they are easy to cultivate, able to grow in high cell densities and can produce high concentrations of lipids. Previously we had isolated a promising lipid producing yeast, *Zygosaccharomyces siamensis* AP1, from wild honey of Central Sulawesi, Indonesia. The strain was able to produce around 19% (w/w biomass) of lipid on glucose. However, its ability to produce lipid on other types of carbon source is still not studied. In this study, we compared the strain ability to produce lipid on glycerol and glucose using different C/N ratio. We also investigated optimum temperature, pH, and growth time for lipid production on glycerol. We found that growing the strain on glycerol with C/N ratio 120 gave the highest lipid content (29.7%). We also found that optimum growth temperature and pH are 30°C and 5, respectively. Maximum lipid biomass, 0.28 g/L, was produced when the strain was grown in the optimum conditions for 72 hours. In conclusion, our results suggest that glycerol is a potential cheap carbon source for lipid production from *Z. siamensis* AP1. We suggest further study to increase lipid yield either by fermentation or strain improvement.

## 1. Introduction

Oleaginous yeasts have been known to produce more than 20% of lipid intracellularly. This capability attracts many researchers to use the fungi as single-cell oil producer to replace oil plants. The oil from yeast, or commonly called Single Cell Oils (SCO), is considered to be suitable as a raw material in biodiesel production [1]. Other researchers consider the oil can be used as a health supplement due to its high PUFA content [2].

Various species of oleaginous yeast have been found; the number is around 5% of known yeast species [2]. The best-known oleaginous yeast genera are *Yarrowia, Candida, Rhodotorula, Rhodosporidium, Cryptococcus, Trichosporon,* and *Lipomyces* [3]. Oleaginous yeast from Indonesia also has been reported. Garay *et al.* [4] reported *Rhodotorula mucilaginosa, Sporidiobolus ruineniae,* and *Sporobolomyces bannaensis* isolated from soil and leaves of Sulawesi able to produce lipid. Other study reported isolating oleaginous *Zygosaccharomyces siamensis* AP1 from wild forest honey of Central Sulawesi [5]. The yeast considered as potential because it can grow in an osmophilic medium allowing the use of a high concentration carbon source to produce lipid.

Many factors influence the production of lipid; one of them is medium composition. Lipid is synthesised in the cytoplasm from acetyl-CoA. The precursor is derived from overproduced citrate excreted by mitochondria as a result of deregulation of the citric acid cycle due to depletion of nitrogen in the medium [2, 6]. Growing oleaginous yeast in medium containing high glucose concentration with minimal nitrogen (C/N ratio >30) will induce the lipid synthesis [7]. Besides glucose, other carbon sources also can be used by yeast cell to produce lipid, such as glycerol and xylose [6].

Glycerol is an abundant side product of biodiesel industry; one kg of glycerol can be obtained from the conversion of 10 kg rapeseed oil into the product. As an industrial waste, glycerol is cheaply available and not many microorganisms able to bioconvert it [8]. Those potencies attract researchers to use glycerol in biodiesel production on many yeast species, such as *Candida curvatus*, *Rhodosporodium toruloides, Yarrowia lipolytica,* and many more [3]. All those studies show the promising result of glycerol. The Indonesian yeast *Z. siamensis* AP1 has shown good quality as oleaginous yeast according to the previous study [5]. However, we do not know if glycerol is a suitable substrate for it.

In this study, we explored *Z. siamensis* AP1 performance in producing lipid using glycerol and glucose as carbon sources. We also observed optimum temperature and pH to grow the strain in glycerol, and finally, we followed the cell growth and lipid production through time. Our goal is to find the best medium composition for the strain and proves that the AP1 can become a potential candidate for SCO industry.

## 2. Material and Methods

### 2.1. Material

*Zygosaccharomyces siamensis* AP1 is a collection of Laboratory of Microbiology, Faculty of Biology, Universitas Gadjah Mada, Indonesia. The strain was kept on Potato Dextrose Agar (Oxoid, Hampshire, UK) slant medium. Glycerol (98%) was obtained from Brataco Chemicals (Jakarta, Indonesia). Glucose, peptone, yeast extract, and malt extract were produced by HiMedia Laboratories (Mumbai, India). Hydrochloric acid (p.a.), sodium hydroxide (p.a.), chloroform, and methanol were obtained from Merck & Co. (Kenilworth, New Jersey, USA), while MgSO_4_.7H_2_O was obtained from BDH (Merck Group, Kenilworth, New Jersey, USA).

### 2.2. Seed culture preparation

Five mL cell suspension of 2-days old AP1 in sterile water was inoculated into 45 mL broth medium containing yeast extract (3 g/L), malt extract (3 g/L), peptone (5 g/L), and glucose (10 g/L). The medium was incubated in a rotary shaker incubator at 30 °C and 220 rpm until the optical density at λ=600nm reached around 0.75 [9].

### 2.3. Determination of substrate C/N ratio for lipid production using glucose and glycerol

Five mL seed culture was inoculated into 45 mL of broth lipid production medium containing 1 g/L yeast extract, 1 g/L peptone, 0.15 g/L MgSO_4_.7H_2_O, and various concentration of glucose or glycerol. The concentrations of the carbon sources were calculated based on Zapata-Vélez and Trujillo-Roldán [10] to meet the defined substrate C/N ratios (Table 1). The inoculated medium incubated in a rotary shaker incubator at 30 °C and 220 rpm for 72 hours. Lipid content from each variation was determined, and the C/N ratio, which gave the value, was used in the next steps. All experiments were done in triplicate.

**Table 1.**
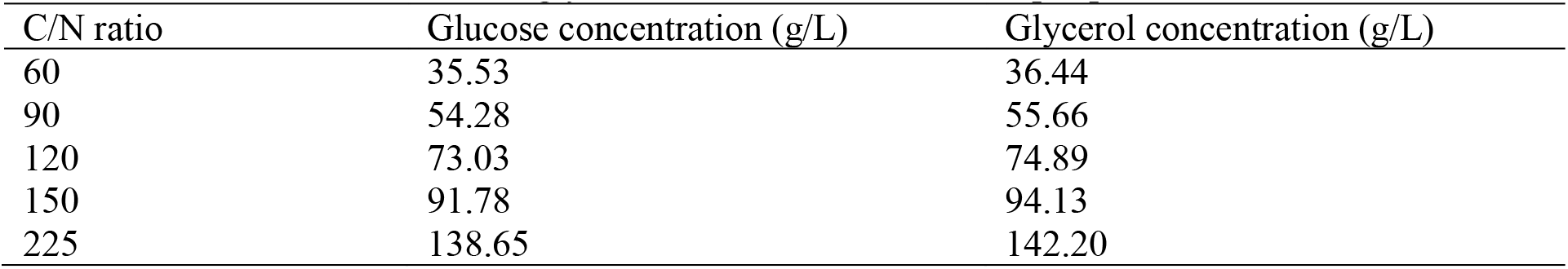
Glucose and glycerol concentrations used in lipid production medium

| C/N ratio | Glucose concentration (g/L) | Glycerol concentration (g/L) |
| --- | --- | --- |
| 60 | 35.53 | 36.44 |
| 90 | 54.28 | 55.66 |
| 120 | 73.03 | 74.89 |
| 150 | 91.78 | 94.13 |
| 225 | 138.65 | 142.20 |

### 2.4. Determination of temperature and initial medium pH for lipid production

The optimum temperature for lipid production was determined by growing AP1 in 45 mL medium containing 1 g/L yeast extract, 1 g/L peptone, 0.15 g/L MgSO_4_.7H_2_O, and carbon source with concentration which gave the best result in the previous experiment. Five mL of seed culture was used. The production was done in a rotary shaker for 72 hours at 220 rpm and various temperatures (25, 30, 35, 40 °C). Lipid biomass from each experiment was calculated. All experiments were done in triplicate.

Initial pH determination was done using medium composition and method similar to the previous step. However, instead of using unadjusted medium pH, we adjusted the pH using hydrochloric acid and sodium hydroxide to the value of 4, 5, 6, and 7 before sterilisation. Temperature value, which gave the best result from the previous step, was used. Lipid biomass from each variation was calculated. All experiments were done in triplicate.

### 2.5. Production profile against time

Production profile was determined by inoculating 5 mL of AP1 seed culture into 45 mL medium containing 1 g/L yeast extract, 1 g/L peptone, 0.15 g/L MgSO_4_.7H_2_O, and carbon source with concentration which gave the best result in the previous experiment. The medium pH was adjusted to the optimum value according to the result of previous experiments. Incubation was done in a rotary shaker at 220 rpm using optimum temperature for five days. The lipid and total biomass were determined daily. All experiments were done in triplicate.

### 2.6. Determination of lipid biomass, total biomass, and lipid content

Lipid biomass was determined according to the method by Bligh & Dyer [11]. A 35 mL of production medium containing yeast cells was centrifuged for 10 minutes at 4000 rotation per minute (rpm). The pellet was collected and washed using 10 mL distilled water twice. On each washing step, water and cells were separated using centrifugation. After that, 10 mL hydrochloric acid 4M was added and the mixture incubated for 2 hours with agitation in room temperature. Next, 5 mL chloroform and 10 mL methanol were added to the mixture, and then it was incubated again for 2 hours in room temperature. After incubation, 5 mL chloroform was added, and the mixture was centrifuged for 10 minutes at 4000 rpm. The organic phase containing lipid at the bottom layer was taken, and all the solvent was evaporated before weighing the remaining lipid. The lipid biomass was expressed as gr/L medium.

Yeast biomass was determined by centrifuging 10 mL of production medium containing yeast cells for 10 minutes at 4000 rpm. The pellet was washed twice as described before drying in an oven at 60°C until constant weight achieved. The biomass was expressed as gr/L medium.

Lipid content is defined as an amount of lipid biomass per total biomass and expressed in percentage (%) [12]. Non-lipid biomass was calculated by subtracting lipid biomass from total biomass.

### 2.7. Data analysis

Graphical presentations of obtained data were made using Microsoft Excel® 2010. The data were analysed statistically using Analysis ToolPak in Microsoft Excel® 2010.

## 3. Results and Discussion

### 3.1. Results

#### 3.1.1. Substrate C/N ratio for lipid production using glucose and glycerol

The effect of C/N ratios to lipid production was observed by measuring lipid content of *Z. siamensis* AP1 after grown for three days on production medium containing glucose or glycerol as carbon source. The results are presented as a clustered column chart (Figure 1).

**Figure 1.**
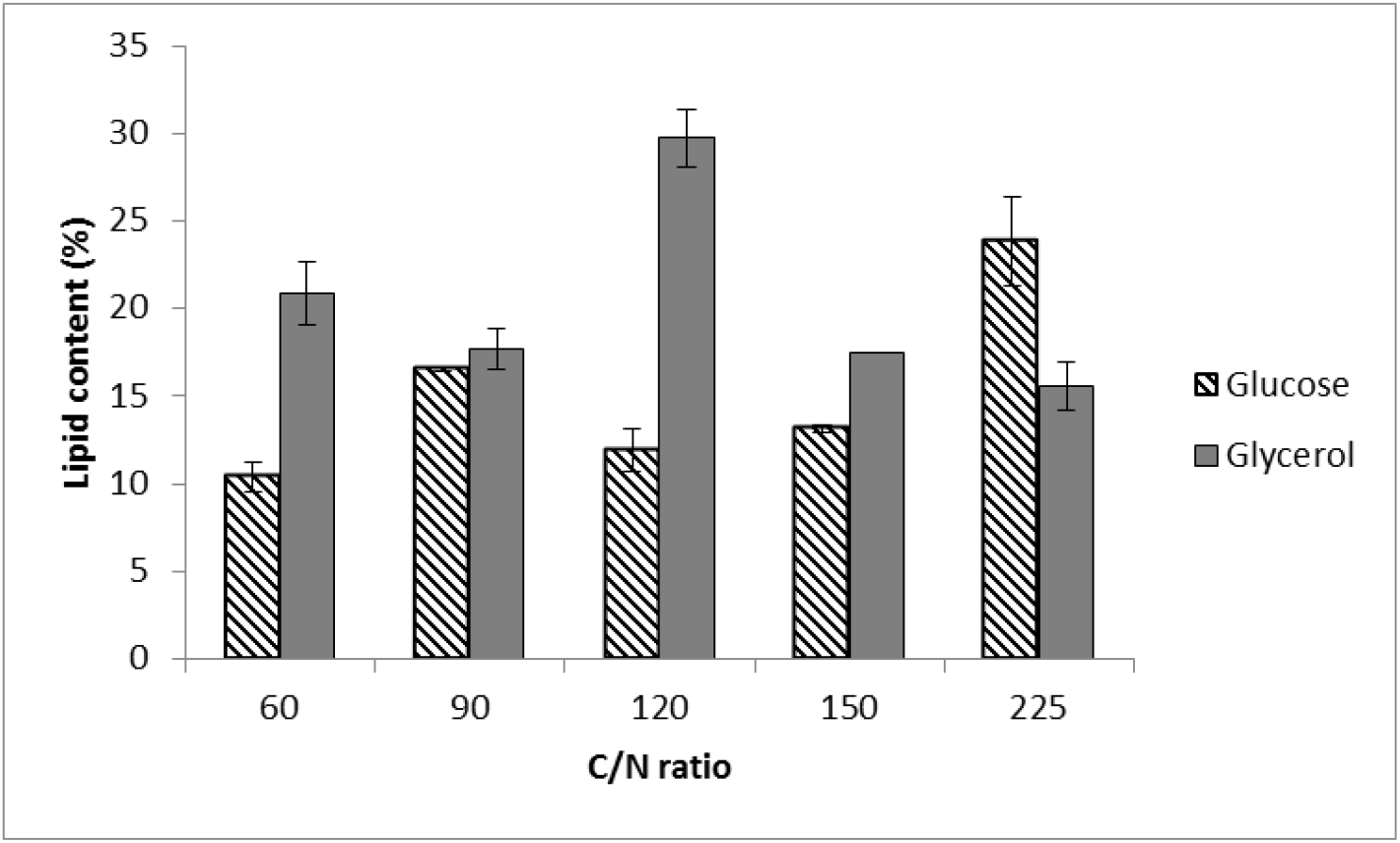
Lipid contents of *Z. siamensis* AP1 after grown on medium with different C/N ratio.

The result shows that *Z. siamensis* AP1 was able to produce higher lipid when was grown on glycerol compared with on glucose for almost all C/N ratios used in the experiment, except for C/N ratio 225. At C/N ratio 120, glycerol gave the most significant higher results than glucose (P=0.0003 at α=0.05), while at C/N ratio 90 the difference was not significant (P=0.436 at α=0.05). Based on the result, glycerol was used as a carbon source for the rest of this study with the medium C/N ratio at 120.

#### 3.1.2. Effect of growth temperature and initial medium pH on lipid production

Effect of growth temperature was determined by growing AP1 in production medium with glycerol as carbon source (C/N ratio 120) at various incubation temperatures for three days. Lipid biomass obtained from the yeast measured, and the result is presented as a column chart (Figure 2).

**Figure 2.**
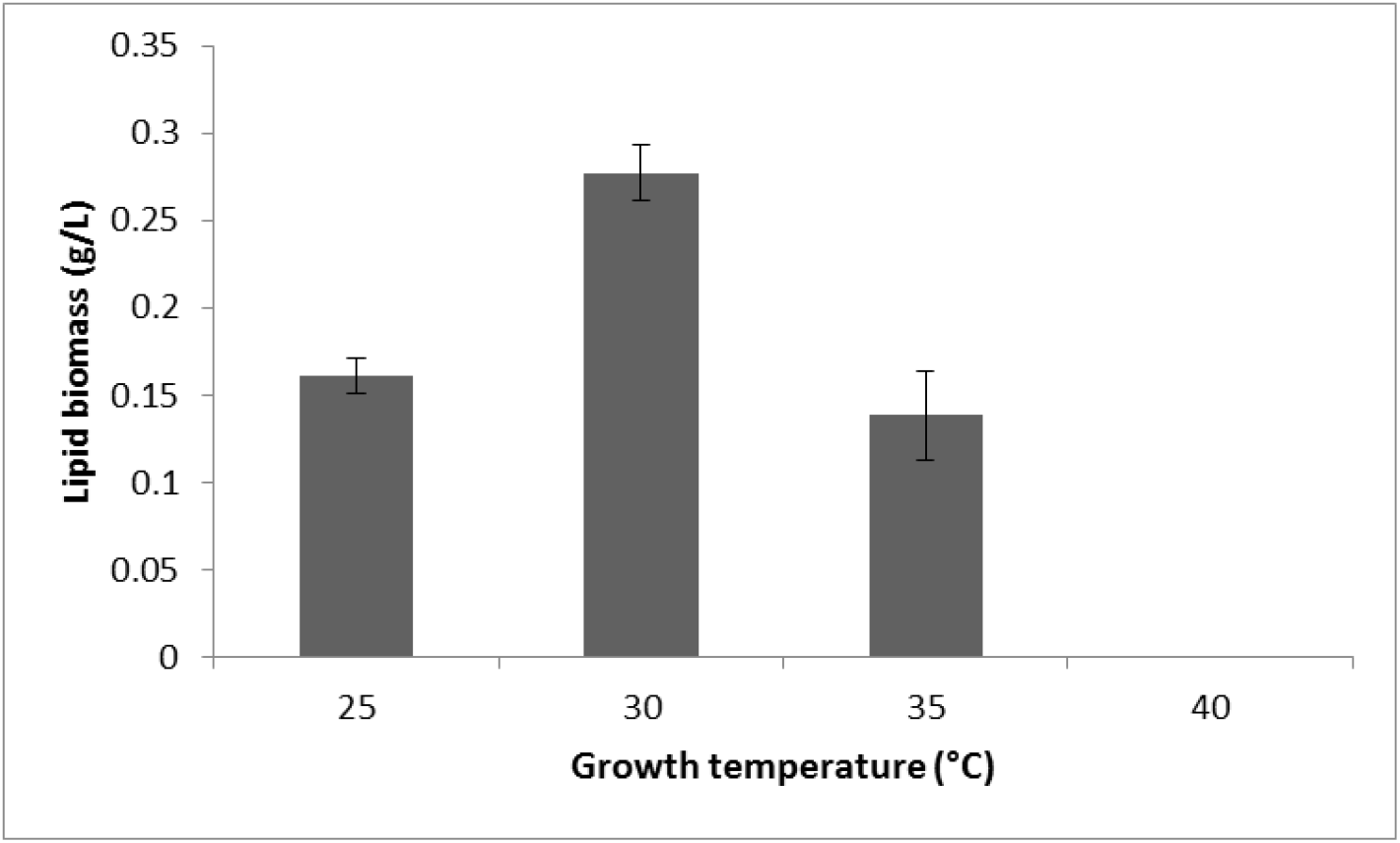
Lipid biomass produced by *Z. siamensis* AP1 in different growth temperatures.

Figure 2 shows that strain AP1 produced the most lipid in growth temperature of 30 °C (P<0.0001 at α=0.05). No growth was observed at temperature 40 °C, indicating the mesophilic nature of the strain. The growth temperature of 30 °C was used in the next experiments based on the result.

Effect of initial pH on lipid production was observed by growing AP1 in production medium as described above using 30 °C as growth temperature. The initial pH of the medium was adjusted to a range of 4 to 7. After three days, lipid biomass was measured, and the result is presented in Figure 3.

**Figure 3.**
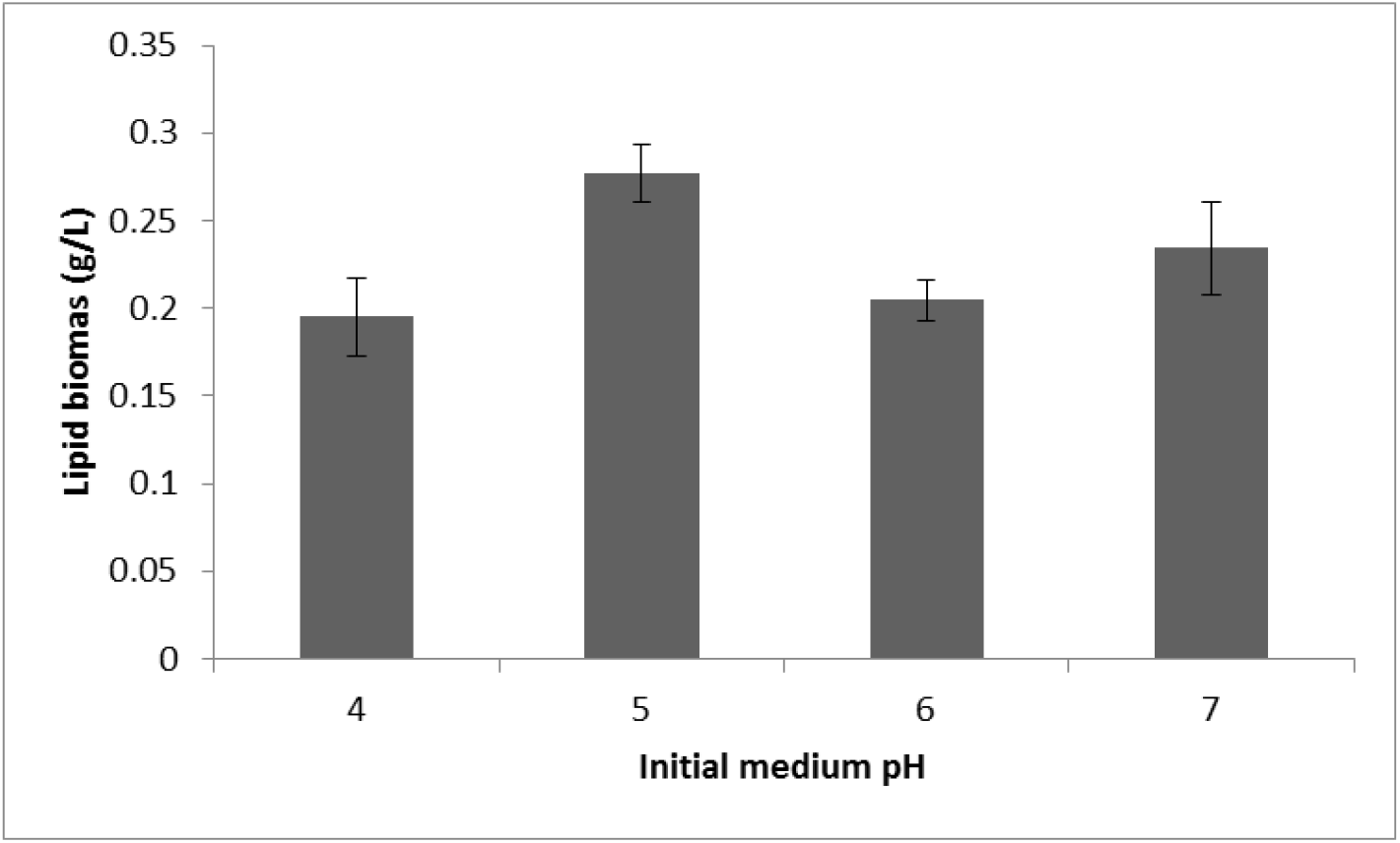
Lipid biomass produced by *Z. siamensis* AP1 in different initial medium pH.

The result showed that strain AP1 produced the highest amount of lipid biomass when it was grown in acidic medium (initial pH 5). The value was significantly higher (P=0.014 at α=0.05) compared with the other results. Based on that, we used pH 5 as the initial medium pH in the next experiments.

#### 3.1.3. Lipid production profile against time

Production profile was determined by growing *Z. siamensis* AP1 for five days in production medium containing optimum composition and using optimum growth condition as discovered at previous experiments. Every day total cell biomass and lipid biomass were measured, then non-lipid biomass calculated by subtracting lipid biomass from total biomass. The result can be seen in Figure 4.

**Figure 4.**
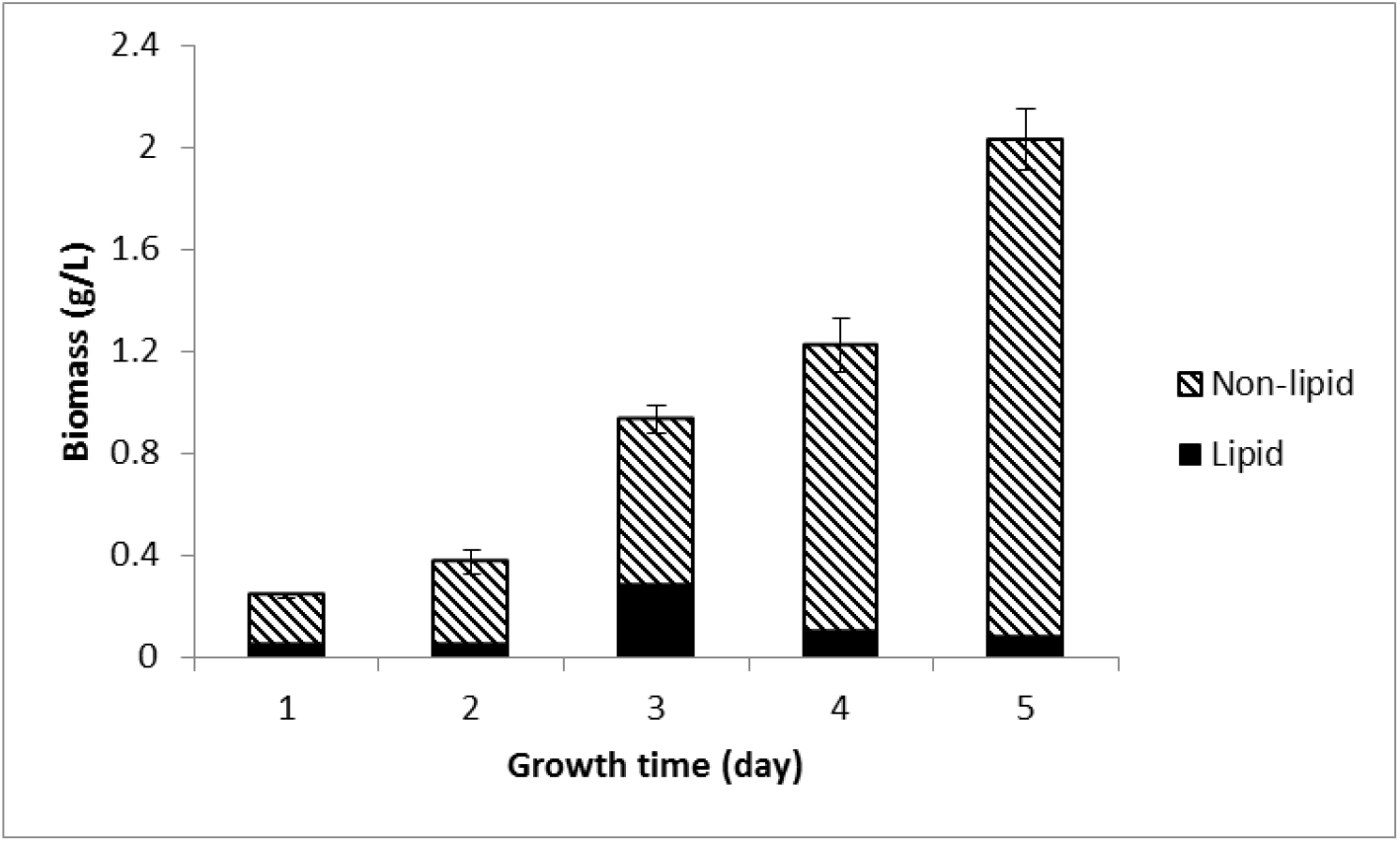
Biomass and lipid production profile of *Z. siamensis* AP1 against growth time.

The growth profile (Figure 4) shows that the total biomass of the strain increased along with time. However, the average of lipid production peaked after three days (0.277 g/L) and decreased afterwards. Like lipid production, lipid content also peaked at day 3 (29.7%) and decreased to a low value on day 5.

### 3.2. Discussion

*Zygosaccharomyces siamensis* AP1 is Indonesian oleaginous yeast, which initially reported able to produce 19% lipid content on 100 g/L glucose (C/N ratio 32) [5]. In order to increase the strain value as potential lipid producer, we optimise its production factors to obtain the highest lipid yield. Although many factors involved in lipid synthesis [13], we only concentrated on carbon source, pH, temperature, and time.

Based on our study, we can increase the AP1 lipid content up to 30% by using 75 g/L glycerol (C/N ratio 120). Glycerol is cheaper than glucose and readily available as industrial waste [8]. Thus, its utilisation for yeast lipid production is growing in the last years [3]. However, glycerol inhibition to the growth of *Candida freyschussii* ATCC 18737 at the concentration higher than 40 g/L has been reported leading to various attempts to prevent it [14]. Our results show that not only the strain AP1 able to produce 30% lipid in 75 g/L glycerol, it also grew well in the medium. The data suggests that the strain resistant to the negative effect of high concentration of glycerol.

In addition to able to use glycerol, *Z. siamensis* AP1 also produced maximum lipid in temperature 30 °C and pH 5. The growth temperature is moderate and put the yeast in the mesophilic zone, while the growth pH indicates the strain is slightly acidophilic. Those growth conditions are tightly related to the yeast natural habitat, which is wild forest honey from Central Sulawesi, which assumed to have a moderate temperature with an acidic condition [15]. The moderate temperature and low pH can be useful if the strain is used at the industrial level; it will not require high energy to grow the strain while the low pH will prevent contamination.

The lipid production profile of *Z. siamensis* AP1 shows us the highest lipid production at day 3 of growth. It assumed that, at day 3, nitrogen in the medium is depleted and cell growth is halted. The remaining glycerol then metabolised through the lipid synthesis pathway. The lipid then degraded via β-oxidation and the cell growth continues [2]. This result also confirms that AP1 growth is not inhibited by high glycerol concentration, as mention above.

Even though strain AP1 has potential futures as described previously, the optimum lipid yield obtained from AP1 is only 0.277 g/L. This value is low if compared to various studies which reported lipid yields higher than 10 g/L [3]. Our finding opens research opportunity to increase the lipid yield of AP1 either by fermentation or strain improvement.

### 4. Conclusions

We concluded that *Z. siamensis* AP1 could use glycerol as a sole carbon source for growth and lipid production. The highest lipid yield, 0.277 g/L, was obtained by growing the yeast in 74.89 g/L at 30°C pH 5 for three days. The yield considered very low compared to reported studies. Hence further studies to increase it are suggested.

## Acknowledgements

The Faculty of Biology, UGM covered laboratory expenses of this study through Lecturer-Student Research Grant 2019 (nr. UGM/BI/1700/M/02/05). Universitas Gadjah Mada covered dissemination and publication expenses of the results of this study through RTA Program 2019 (nr. 3002/UN1/DITLIT/DIT-LIT/LT/2019).

